# The Resting-State Causal Human Connectome is Characterized by Hub Connectivity of Executive and Attentional Networks

**DOI:** 10.1101/2021.10.20.465211

**Authors:** Eric Rawls, Erich Kummerfeld, Bryon A. Mueller, Sisi Ma, Anna Zilverstand

## Abstract

We demonstrate a data-driven approach for calculating a “causal connectome” of directed connectivity from resting-state fMRI data using a greedy adjacency search and pairwise non-Gaussian edge orientations. We used this approach to construct n=442 causal connectomes. These connectomes were very sparse in comparison to typical Pearson correlation-based graphs (roughly 2.25% edge density) yet were fully connected in nearly all cases. Prominent highly connected hubs of the causal connectome were situated in attentional (dorsal attention) and executive (frontoparietal and cingulo-opercular) networks. These hub networks had distinctly different connectivity profiles: attentional networks shared incoming connections with sensory regions and outgoing connections with higher cognitive networks, while executive networks primarily connected to other higher cognitive networks and had a high degree of bidirected connectivity. Virtual lesion analyses accentuated these findings, demonstrating that attentional and executive hub networks are points of critical vulnerability in the human causal connectome. These data highlight the central role of attention and executive control networks in the human cortical connectome and set the stage for future applications of data-driven causal connectivity analysis in psychiatry.

## 1 Introduction

Brain network interactions give rise to information processing and cognition (Bressler, 1995; Bullmore and Sporns, 2009; Friston, 2002; McIntosh, 2000). Brain networks have a non-random topological organization (Bullmore and Sporns, 2009), including both segregated modules and a small number of highly connected nodes (Achard et al., 2006; Eguíluz et al., 2005; Sporns et al., 2007; van den Heuvel and Sporns, 2013), the “hubs” of the connectome. These hubs coordinate the transfer of large amounts of information through brain circuits (Mišić et al., 2015, 2014; van den Heuvel et al., 2012) and play a critical role in coordinating communication between disparate brain regions (Cole et al., 2013; Sporns, 2013, 2012; van den Heuvel and Sporns, 2011). Here we present data on a “causal connectome” derived from resting-state neuroimaging data using a data-driven causal discovery method that first estimates directly connected brain regions without the false positives produced by typical connectivity methods (Reid et al., 2019), then additionally estimates the direction of those connections. We describe the characteristics of this causal connectome, with a specific focus on the central hubs of the resting-state causal connectome. While networks with directional information have several designations in the literature (effective, causal, directed), for consistency we will refer to these networks as causal networks throughout this manuscript.

Hub-like connectivity can be characterized using measures of centrality, a set of metrics that quantify the capacity of a node in a graph to influence (or be influenced by) other nodes. Over 100 measures of centrality have been proposed (Jalili et al., 2015), only a subset of which are commonly applied to brain network analysis (Rubinov and Sporns, 2010; van den Heuvel and Sporns, 2013). The most common is degree centrality (the number of connections attached to each node), which has a simple and intuitive interpretation, but alone provides limited information. For example, a node might have relatively few connections, but still act as an important bottleneck for communication between many other nodes in the network. Such a node could be thought of as a city highway. While it might have relatively few direct connections (entrances and exits), the highway still serves as the quickest path between many locations in the city. To capture this type of connectivity, measures such as betweenness centrality (how often a node lies on the shortest path between two other nodes; Freeman, 1977) have been developed.

Different metrics for centrality are often highly correlated with each other (Oldham et al., 2019; Oldham and Fornito, 2019), but some centrality metrics may be more appropriate for certain types of networks than for others. For example, while degree centrality is frequently used for characterizing structural brain hubs (Crossley et al., 2014; Rubinov et al., 2015), degree in correlation-based fMRI networks is biased to identify nodes that are part of the largest (number of nodes/regions) resting-state networks (RSNs) of the brain (Power et al., 2013; van den Heuvel and Sporns, 2013). Thus, early degree-based analyses often identified high-degree nodes in the default mode network (Buckner et al., 2009; Cole et al., 2010; Power et al., 2013; Tomasi and Volkow, 2011; van den Heuvel and Sporns, 2013), one of the brain’s largest resting-state subnetworks. Because of this confound, groups using Pearson correlation-based network analyses of fMRI data have considered alternative methods for identifying functional hubs, including metrics that consider the diversity of between-RSN connections such as participation coefficient (Bertolero et al., 2018; Grayson et al., 2014; Power et al., 2013; Reber et al., 2021), and coactivation/connectivity over multiple cognitive tasks (Cocuzza et al., 2020; Cole et al., 2013; Crossley et al., 2014, 2013; Ray et al., 2020). These more recent analyses reveal a set of brain hubs distributed broadly through frontal, parietal, and temporal cortices. Notably, defining what constitutes “high” centrality is dependent on the distribution of centralities exhibited by the network, and there is no evidence that one-size-fits-all cutoffs for categorizing network nodes as hubs can be meaningfully applied to brain networks. For example, when a cutoff for defining hubs as proposed in (Guimerà and Nunes Amaral, 2005) was applied to the brain network examined in Power et al. (2013), only a single local hub was identified. As such, the distinction between what constitutes a hub as opposed to a non-hub is essentially arbitrary (Oldham and Fornito, 2019). Our analysis of hub connectivity will thus focus on continuous measures of centrality, providing a quantitative comparative measure of which brain networks or nodes are more hub-like than others in the investigated causal connectome.

Critical to our current investigation, most resting-state fMRI connectivity studies use Pearson correlations to estimate undirected connectomes, providing information about connected brain regions (hereafter adjacencies), but not the direction of these connections (hereafter orientations). There is a recognized need for network modeling methods that can accurately estimate adjacencies without the false positives inherent to correlation-based approaches (Reid et al., 2019; Smith et al., 2011), as well as estimating the direction or orientation of these edges (referred to as causal or effective connectivity (Ramsey et al., 2010; Reid et al., 2019; Smith, 2012). However, current methods for fMRI resting-state causal connectivity are limited (Ramsey et al., 2014, 2010; Sanchez-Romero et al., 2019; Smith et al., 2011). Granger causality (Granger, 1969) attempts to recover causal influences using time-lagged regressions. Smith et al. (2011) found that several variations of Granger causality have negligible accuracy in detecting adjacencies or orientations in simulated fMRI data, and Sanchez-Romero et al. (2019) tested two additional variations of Granger causality (multivariate Granger causality; Barnett and Seth, 2014; autoregressive modeling with permutation testing; Gilson et al., 2017), finding that these more recent methods also had low precision for adjacencies and orientations in the presence of realistic noise. GIMME (Gates and Molenaar, 2012), a group-level algorithm that also uses time lags to infer causality, achieves better performance, but is computationally intensive and can only scale to a small number of brain regions (Sanchez-Romero et al., 2019). Dynamic causal modeling (DCM; (Friston et al., 2003) was originally designed for task-based fMRI data, but the stochastic DCM (Li et al., 2011) and spectral DCM (Friston et al., 2014) variants can be applied to resting-state fMRI data (albeit only for a small number of brain regions due to computational demands). Frässle et al. (2021) recently proposed a highly scalable variant of regression DCM (Frässle et al., 2017) for resting-state fMRI data that is scalable enough to support whole-cortex analyses. While the connectomes generated by this newly proposed method appear to have face validity (Frässle et al., 2021), a quantitative analysis of large-scale resting-state connectivity patterns using regression-DCM has not yet been completed. Instead, the current report focuses on a distinct set of methods grounded on Bayes networks, which have promise for uncovering causal connectivity from fMRI (Mumford and Ramsey, 2014; Ramsey et al., 2014; Sanchez-Romero et al., 2019), and for dealing with the high dimensionality of whole-cortex data (Ramsey et al., 2017). Unlike GIMME, spectral DCM, and stochastic DCM, Bayes net methods are highly scalable, and unlike regression DCM, these methods do not require specification of priors or hemodynamic response function and make no assumptions about the physiology giving rise to the observed hemodynamic signal. Suitable combinations of causal discovery-based methods can achieve near-perfect precision and recall in simulated fMRI data (Hyvärinen and Smith, 2013; Ramsey et al., 2014), even for networks with feedback cycles (Sanchez-Romero et al., 2019).

In the current study, we capitalize on these recent advances in causal discovery machine learning to build whole-cortex causal connectomes from single-subject resting-state fMRI data. We apply a variation of a previously proposed (Ramsey et al., 2011; Ramsey et al., 2014; Sanchez-Romero et al., 2019) two-step causal discovery framework that breaks the connectome computation into separate adjacency and orientation steps. For convenience we refer to our approach as GANGO (Greedy Adjacencies and Non-Gaussian Orientations). GANGO first estimates whole-cortex adjacencies using Fast Greedy Equivalence Search (FGES; Ramsey et al., 2017). FGES is a parallelized version of Greedy Equivalence Search (Chickering, 2002), a Bayes Network method with high sensitivity for detecting adjacencies, but poor accuracy for orientations in simulated fMRI data (Smith et al., 2011). Thus, we follow this initial adjacency search with a pairwise edge orientation algorithm that exploits non-Gaussian information in the hemodynamic signal (Hyvärinen and Smith, 2013), shown in (Ramsey et al., 2014; Sanchez-Romero et al., 2019) to have high precision and recall for determining edge orientation of the Smith and colleagues (2011) simulations. We thus obtain, on a single-subject basis, a whole-cortex graph summarizing dominant causal connectivity between brain regions. We focus our investigation on the hub structure of this novel causal connectome. Overall, we demonstrate hub-like causal connectivity profiles of the dorsal attention network, frontoparietal network, and cingulo-opercular network.

## 2 Methods

### 2.1 Subjects

All analyses used publicly available resting-state functional neuroimaging data from 442 unrelated healthy young adult subjects recruited as part of the Washington University – Minnesota (WU-Minn) Human Connectome Project Consortium (56% [n = 248] female; aged 22-35 [mean age = 28.6 years]; https://db.humanconnectome.org/data/projects/HCP_1200) (Barch et al., 2013; Glasser et al., 2013; Marcus et al., 2013; Smith et al., 2013; Uğurbil et al., 2013; Van Essen et al., 2013). All subjects provided written informed consent at Washington University.

### 2.2 Resting-State fMRI Acquisition and Preprocessing

Structural (T1 and T2 images, required for preprocessing functional neuroimaging data) and functional MRI data were collected at Washington University on the Siemens 3T Connectome Skyra scanner. Full details of the acquisition parameters for the HCP data are described in (Uğurbil et al., 2013). Each subject’s resting-state data was collected over two days in four sessions (14:33/session; 1200 samples/session). In this study we analyzed only one day of data (two runs, individually z-scored and concatenated) to limit potential state influences on fMRI measures. Structural and functional data preprocessing is described in (Glasser et al., 2013), and used version 3.21 of the HCP preprocessing pipeline. Structural data preprocessing consisted of bias field and gradient distortion correction, coregistration of T1/T2 images, and registration to MNI space. Cortical surface meshes were constructed using FreeSurfer, transformed to MNI space, registered to individual surfaces, and downsampled. Functional MRI preprocessing consisted of gradient distortion correction, motion correction, EPI distortion correction, followed by T1 registration. Transforms were concatenated and run in a single nonlinear resampling to MNI space followed by intensity normalization. Data were masked by the FreeSurfer brain mask, and volumetric data were mapped to a combined cortical surface vertex and subcortical voxel space (“grayordinates”) using a multimodal surface registration algorithm (Robinson et al., 2014) and smoothed with a 2mm FWHM Gaussian in surface space (thus avoiding smoothing over gyral banks). fMRI data were conservatively high-pass filtered with FWHM = 2000 s and cleaned of artifacts using ICA-FIX (Griffanti et al., 2014; Salimi-Khorshidi et al., 2014). This filter was implemented as a weighted linear function in FSL v6.0.2, which was shown in (Ramsey et al., 2014) to not introduce Gaussian trends into the data (unlike e.g., Butterworth filters or the built-in SPM filter). Artifact components and 24 motion parameters were regressed out of the functional data in a single step, producing the final ICA-FIX denoised version of the data in CIFTI (“grayordinates”) space (Glasser et al., 2016b) that was used in subsequent analyses.

### 2.3 Construction of Whole-Brain Causal Connectomes

Our analysis pipeline began with n = 442 sets of preprocessed, multi-modally surface registered, ICA-FIX denoised fMRI data provided by the HCP consortium. We parcellated cortex surface vertices into 360 regions using a recently developed multimodal parcellation (Glasser et al., 2016a).

We implement a computational strategy to define causal connectomes on a per-subject basis using a two-step process we refer to as GANGO (Greedy Adjacencies and Non-Gaussian Orientations). This approach is motivated by previous work (Ramsey et al., 2011; Ramsey et al., 2014; Sanchez-Romero et al., 2019; Smith et al., 2011) indicating that 1) Bayes net algorithms such as PC (Spirtes et al., 2001) and Greedy Equivalence Search (Chickering, 2002) provide a highly precise solution to identify nodal adjacencies (but not orientations) in simulated fMRI data, and 2) pairwise orientation algorithms based on data skewness can accurately identify edge orientations in simulated fMRI data. In the first step, the GANGO approach defines nodal adjacencies (connected regions) using Fast Greedy Equivalence Search (FGES; Ramsey et al., 2017), a parallelized and highly scalable version of GES. This algorithm finds a sparse set of directed and undirected connections between continuous variables by minimizing a penalized likelihood score over the entire graph, typically scored using the Bayesian Information Criterion (BIC; Schwarz, 1978). FGES proceeds in two stages, first adding edges until the BIC stops improving, then removing edges until the BIC stops improving (Ramsey et al., 2017). While FGES has not commonly been used for analysis of empirical neuroimaging data, this method was applied (in combination with direct stimulation) to test causal connectivity patterns of the amygdala (Dubois et al., 2020) with promising initial findings for mapping human emotion networks. We computed FGES with causal-cmd v1.2.0 (https://bd2kccd.github.io/docs/causal-cmd/) using default parameters (BIC penalized likelihood score, penalty discount = 1 corresponding to the classic BIC score). GES has been shown in simulations to obtain highly accurate estimates of nodal adjacencies, but relatively inaccurate orientations (Smith et al., 2011). Therefore, we made the FGES-derived graph undirected by symmetrizing it across the diagonal.

The GANGO approach then orients these undirected edges using non-Gaussian information in the BOLD signal. We applied an estimate of the direction of causal effect based on pairwise likelihood ratios under the linear non-Gaussian acyclic model (Hyvärinen and Smith, 2013). Several approaches have been proposed to orient causal graph edges using non-Gaussian information. For example, Ramsey et al. (2011) used IMaGES (a group-level version of GES) to infer adjacencies from fMRI data and proposed two early measures for orienting edges using non-Gaussian information. More recent approaches (Hyvärinen and Smith, 2013; Ramsey et al., 2014; Sanchez-Romero et al., 2019) have built on these early algorithms, with improved orientation accuracy. For the GANGO framework, we adopt the RSkew method, an outlier-robust skew-based measure (Hyvärinen and Smith, 2013). RSkew has shown to generate optimal estimates of causal direction in simulated fMRI data (Ramsey et al., 2014; Sanchez-Romero et al., 2019), and was calculated using the authors’ MATLAB implementation of RSkew (https://www.cs.helsinki.fi/u/ahyvarin/code/pwcausal/; RSkew is method 4). Hyvärinen and Smith (2013) provide an explanation of how non-Gaussian information can be used to orient edges between pairs of variables. Assuming x -> y, both variables will have large values in cases where x is large (but x will not necessarily take on large values when y is large). Due to regression towards the mean, the value of x must typically be larger than that of y. Cumulant-based approaches such as RSkew, the method used here, calculate pairwise contrasts that magnify extreme values of either x or y, allowing determination of the most likely causal direction. For an in-depth explanation, we refer the reader to (Hyvärinen and Smith, 2013).

Since the RSkew orientation method requires that data be skewed to obtain accurate measures, we tested whether the resting-state data met these assumptions. For each subject, we tested whether the BOLD data were significantly skewed with reference to Gaussian data using an approach adapted from Sanchez-Romero et al. (2019). Within each single participant, we calculated the skewness of the BOLD time series separately for each parcel, resulting in 360 skewness values per subject. For each subject, we then simulated 360 Gaussian time series of the same length as the BOLD data (n = 2400 points) for use as surrogate data and calculated the skewness of each of the Gaussian time series. For each subject, we then statistically tested whether the skewness of the observed BOLD data (n = 360 values) exceeded the skewness derived from Gaussian surrogate data (n = 360 values) using a one-tailed Wilcoxon rank sum test. Since both positive and negative skewness values indicate skewed data, for all analyses we used the absolute value of skewness.

### 2.4 Resting-State Network Connectivity Statistics

For each subject, we categorized brain regions into 12 resting-state networks (RSNs) using the recently developed Cole-Anticevic Brain-wide Network Partition (CAB-NP) (Ji et al., 2019). We established whether each RSN shared a statistically significant proportion of connections with each other RSN (*Figure 1b*) by calculating the total number of connections that the RSN shared with the other 11 RSNs (the total number of out-of-RSN connections). Then, by dividing that total by 11, we arrive at a suitable null hypothesis of equal inter-RSN connectivity (i.e., that the RSN shared out-of-RSN connections equally between the other 11 RSNs). For each of the other 11 RSNs, we tested the actual number of shared between-RSN connections against the null hypothesis of equal connectivity to all 11 RSNs using Wilcoxon signed-rank tests, which we corrected for multiple corrections using the false discovery rate (FDR) procedure (Benjamini and Hochberg, 1995), thus establishing whether pairs of RSNs were significantly connected. To clarify the direction of causal connectivity between significantly connected pairs of RSNs, we calculated the proportion of causal connections from each RSN to each other RSN, then compared that proportion against a null hypothesis of 50% (i.e., that the connections between the pair of RSNs A & B are equally from A to B and from B to A) using Wilcoxon signed-rank tests (FDR-corrected for multiple comparisons). For each pair of RSNs found to be significant with reference to a null hypothesis of equal (i.e., random) inter-RSN connectivity, we calculated the effect size of the test for significant shared connections using Cohen’s d, and masked inter-RSN connections that did not meet at least the threshold for a small effect size (Cohen’s d >= 0.2). We plot this result in the form of a force-directed graph, by first drawing connections between pairs of significantly connected networks. If we were able to determine a significant direction of connectivity, this connection was unidirectional; otherwise, the connection was drawn bidirectional. For significantly one-directional connections, we also recorded the proportion of connections going in the significant direction. This analysis clarified a) which RSNs are significantly connected with at least a small effect size, and b) which RSNs significantly send (vs. receive) information to (vs. from) other RSNs. We supplemented this with an analysis of participation coefficient (a measure of the diversity of RSNs a node connects to), calculated separately for incoming and outgoing connections. Participation coefficient *P*_*i*_ of node i is calculated according to the equation

**Figure 1.**
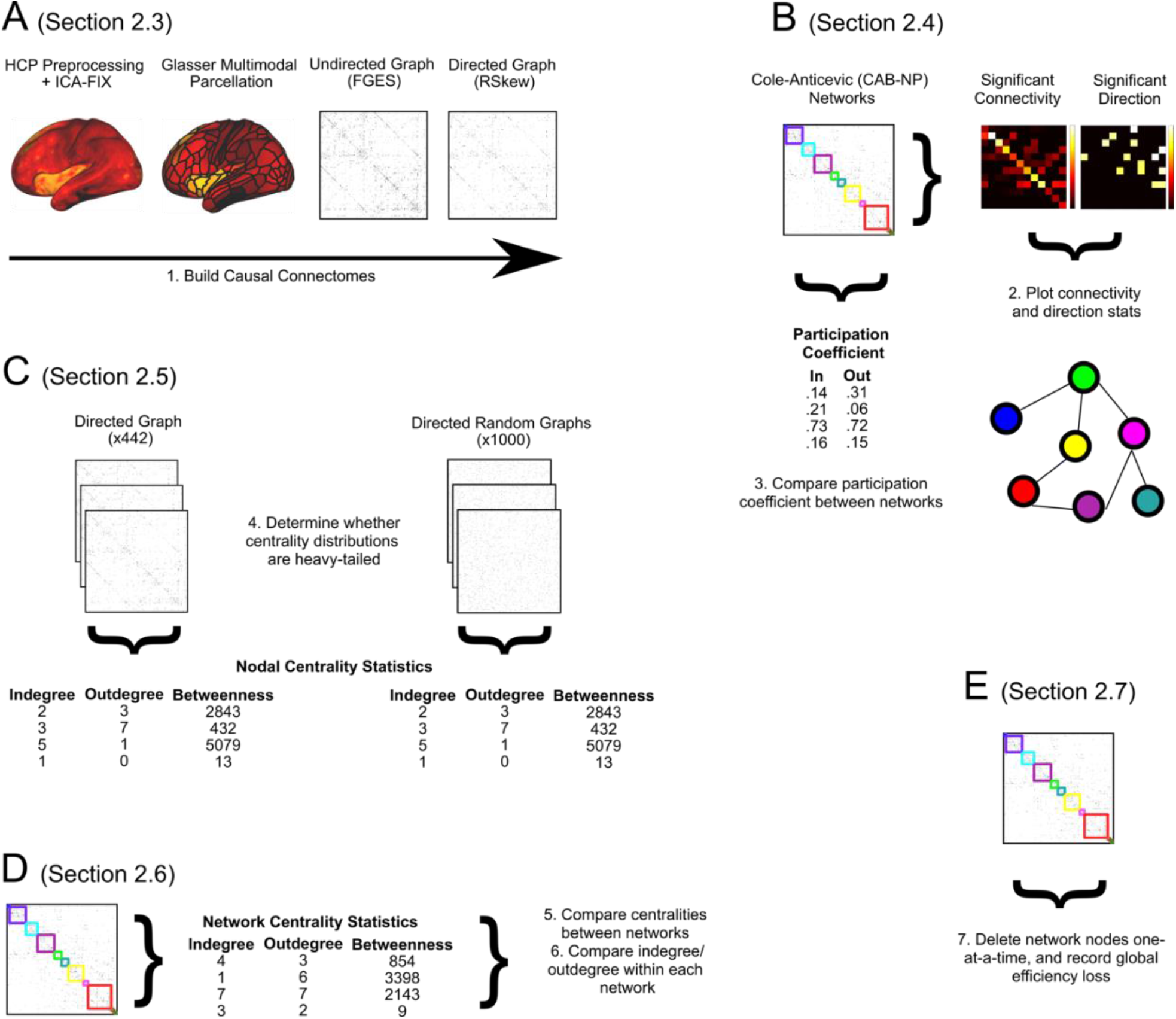
Summary of the strategy we employed to build single-subject (n = 442) causal connectivity graphs for further analysis, and the analyses we ran to characterize the hub structure of these RSNs. **A**: Described in Section 2.3 (Methods). The brain surface in the plot is a single TR of a randomly selected subject’s resting-state data, representing the resting-state activation maps from which causal connectomes were computed, to illustrate the preprocessing steps. **B**: Described in Section 2.4 (Methods) and Section 3.3 (Results). **C**: Described in Section 2.5 (Methods) and Section 3.4 (Results). **D**: Described in Section 2.6 (Methods) and Sections 3.5 and 3.6 (Results). **E**: Described in Section 2.7 (Methods) and Section 3.7 (Results).

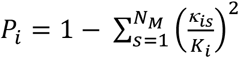 where *Κ*_*is*_ is the number of connections between node *i* and RSN *s* and *K*_*i*_ is the total degree of node *i* (Guimerà and Nunes Amaral, 2005). For nodes that connect entirely within their own RSN, this equation results in a participation coefficient of zero, and for nodes that connect homogeneously over all RSNs the participation coefficient will approach one (Guimerà and Nunes Amaral, 2005; Power et al., 2013). We computed participation coefficients using the *part_coeff* function in the Brain Connectivity Toolbox (Rubinov and Sporns, 2010). We compared nodal participation coefficients in 12 established RSNs (Ji et al., 2019) that have been validated for our current parcellation using Friedman tests (since network is a within-subject measure) and conducted post-hoc pairwise comparisons with control for multiple comparisons using the Nemenyi test.

### 2.5 Centrality Distributions in Human Cortex

For each causal graph, we calculated nodal (n = 360) centrality statistics using indegree (number of incoming connections), outdegree (number of outgoing connections), and betweenness centrality (number of shortest paths the node participates in) (*Figure 1c*). To statistically quantify whether centrality-based cortical hubs existed based on significantly heavy-tailed centrality distributions, we generated a reference set of 1000 random directed graphs with the same number of nodes (360) and connections as the cortical causal graphs (*Figure 1c*). Specifically, for each run (of 1000), we chose the exact number of connections by drawing a random number from a normal distribution with the same mean and standard deviation as the number of connections across subjects (mean n connections = 1452, SD = 107); thus, the surrogate graphs approximated the distribution of connection counts present in the actual data. Random graphs were created using the *makerandCIJ_dir* function from the Brain Connectivity Toolbox (Rubinov and Sporns, 2010), which creates a random directed (causal) graph with a specified number of connections. As the degree distribution is the main variable in this analysis, this function (adding connections randomly off the diagonal) produces random graphs with no constraints other than having the overall number of connections be equivalent to the observed data. This does not bias the resulting graphs to have small-world properties (e.g. an Erdős–Rényi graph; Erdős & Rényi, 1960) or to be constrained to having the same degree distribution as the FGES graphs (e.g. randomizing graphs derived from FGES). We then used Wilcoxon rank-sum tests to compare the skewness of centrality distributions (indegree, outdegree, betweenness) of the causal connectivity graphs to the random graphs. We additionally applied this permutation analysis to categorize cortical nodes as hubs or non-hubs, based on whether a node’s centrality exceeded the 95^th^ percentile of the surrogate distribution (i.e., whether a node was significantly in the highly central tail of the distribution). These binary masks of regions thus categorized as significant hubs are provided in the supplement.

### 2.6 Resting-State Network Differences in Centrality-Based Hubs

We compared the average nodal centralities (degree, betweenness) in 12 established RSNs (Ji et al., 2019) that have been validated for our current parcellation (*Figure 1d*), to obtain a continuous ranking of which networks are the most “hub-like” in the causal connectome. For each subject, we calculated the average nodal indegree, outdegree, total degree, and betweenness centrality for nodes within each of these 12 RSNs using the MATLAB *centrality* function. We compared centrality across the 12 RSNs using Friedman tests and conducted post-hoc pairwise comparisons with control for multiple comparisons using Nemenyi tests. Additionally, we compared indegree and outdegree within each of the 12 RSNs using Wilcoxon signed-rank tests, and FDR-corrected the resulting *p*-values for n = 12 multiple comparisons.

### 2.7 Network Vulnerability to Targeted and Random Attack

To clarify the functional role of hubs in the causal cortical network, we subjected each subject’s causal connectome to a targeted attack analysis. Broadly, virtual attack analyses proceed by deleting nodes from the network and recording some measure of whole-network fitness as nodes are removed. The dependent value in this analysis is chosen to be a measure of network functional integration, to characterize nodes that are most critical for network communication (Rubinov and Sporns, 2010). As discussed in (Rubinov and Sporns, 2010), the most common measure of network functional integration is the characteristic path length (Watts and Strogatz, 1998). However, characteristic path length cannot be meaningfully computed on networks with disconnected nodes, as disconnected nodes are defined to have infinite path length. Thus, authors have argued that global efficiency (the inverse shortest path lengths in the network, defined to be zero for disconnected nodes (Latora and Marchiori, 2001)) is a superior measure of network functional integration (Achard and Bullmore, 2007; Bassett and Bullmore, 2006; Rubinov and Sporns, 2010). As such, network global efficiency is typically used as the dependent value in published virtual attack analyses (Crossley et al., 2014; Lin et al., 2018; Lo et al., 2015; van den Heuvel et al., 2018; van den Heuvel and Sporns, 2011).

To assess cortical vulnerability at the RSN level, we deleted the nodes of each RSN (one at a time) from the individual subject-level cortical causal connectivity graphs, and we recorded changes in connectome communication efficiency as percent change in global efficiency (*Figure 1e*), calculated using the *efficiency_bin* function in the Brain Connectivity Toolbox (Rubinov and Sporns, 2010). We supplemented this targeted attack analysis with a random attack analysis, by deleting randomly selected nodes rather than specifically targeting nodes from one RSN at a time. To minimize order effects in the virtual lesion analysis we ran the analysis 10 times for every subject, deleting the nodes within each RSN in a random order each time, then we took the mean of the efficiency loss over the 10 runs. This resulted in a set of 13 loss-of-efficiency curves per subject that quantified how strongly communication was impaired as successive nodes from each RSN were deleted. We took the pointwise derivative of each subject’s loss-of-efficiency curves (for each of 13 deletion schedules) and compared the average pointwise slope between RSNs (plus random deletion) to quantify which RSN resulted in the most rapid loss-of-efficiency. RSN deletion slopes were statistically compared using a Friedman test with post-hoc significance testing using the Nemenyi test. Additionally, we examined which nodes had the greatest overall impact on connectome loss-of-efficiency by deleting (in single subjects) each node (of 360) one at a time and recording the change in overall global efficiency.

## 3 Results

### 3.1 Individual Subjects Functional Neuroimaging Data are non-Gaussian and Skewed

Within each single subject, we compared the skewness of the fMRI time series to the skewness of random Gaussian data. This analysis revealed that every single subject’s BOLD time series were significantly (rank sum tests, *p* < .05) more skewed than surrogate Gaussian data. At the group level, data were significantly more skewed than random Gaussian data as well (*p* < .001). Thus, we conclude that the functional neuroimaging data analyzed in the current report are non-Gaussian, specifically skewed, and therefore it is appropriate to apply the skew-based orientation step to the analysis.

### 3.2 Whole-Cortex Causal Graphs are Sparse but Well-Connected

The resulting whole-cortex causal connectomes were sparse, containing only ∼2.25% of all possible connections (360 parcels: mean n connections = 1452). Nevertheless, the graphs were well-connected – in most graphs (93.7%), every node was connected to at least one other node, and of the 6.3% of graphs that contained disconnected nodes, each graph had very few disconnected nodes (median = 1, max = 2). Thus, despite the sparsity of the cortical causal connectivity graphs, these graphs appear to capture global causal patterns of connectivity.

### 3.3 Diversity of Inter-Network Connections Highlights Hub Roles of Multiple Brain Networks

To summarize the overall connectivity structure of our cortical causal graphs, we categorized each of 360 cortical nodes into 12 resting-state networks (RSNs) (Ji et al., 2019). For a visual of these 12 RSNs plotted on the cortical surface, see Figure 2a. We first examined patterns of connectivity between these large-scale brain RSNs. We established whether each RSN shared a statistically significant proportion of connections with each other RSN. We additionally tested for a preferred direction of connectivity between significantly connected pairs of RSNs.

**Figure 2.**
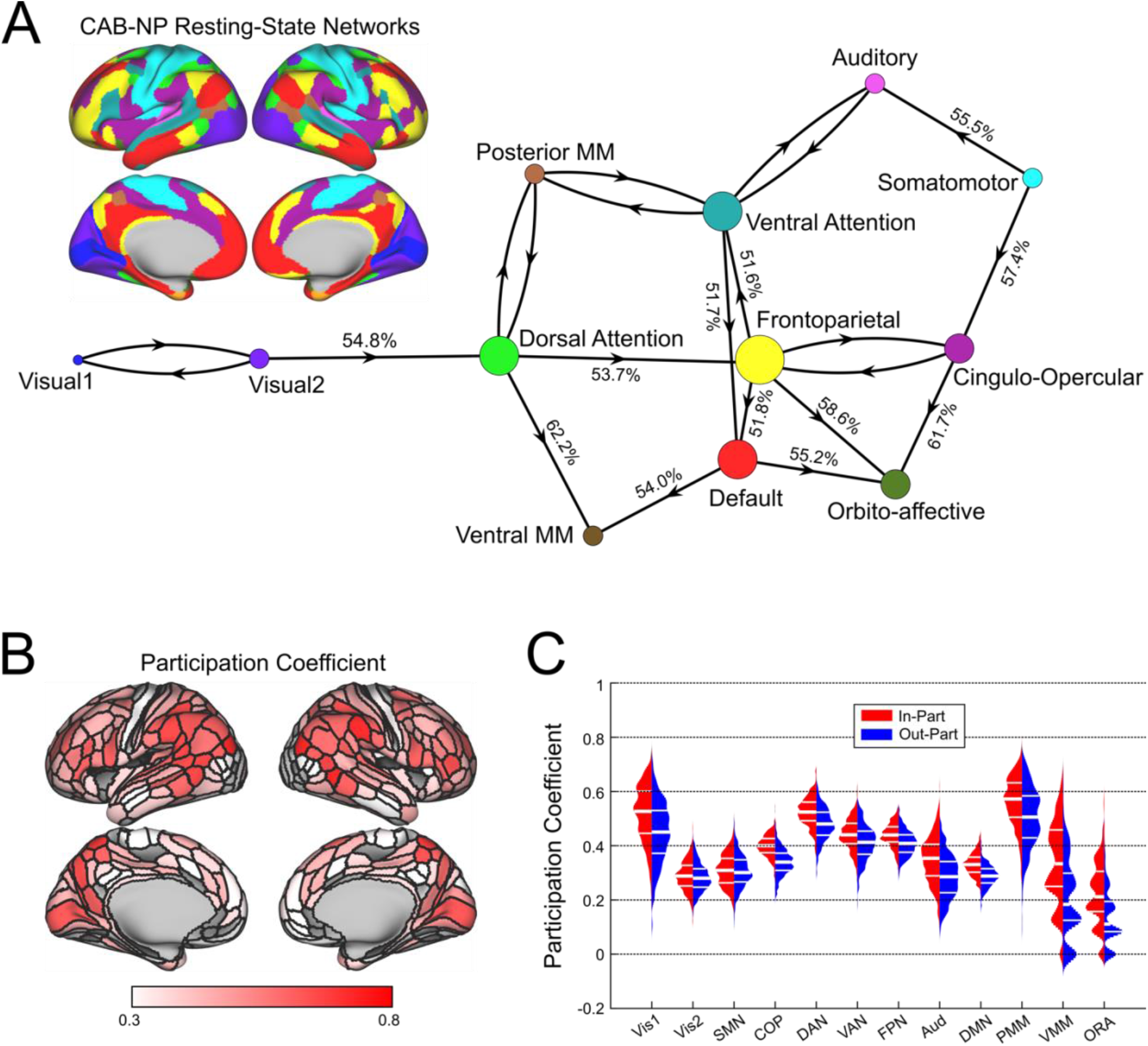
Diversity of Inter-Network Connections Highlights Hub Roles of Multiple Brain Networks. A: The cortical surface is a reference plot of the 12 RSNs from the Cole-Anticevic Brain-Wide Network Partition (CAB-NP; color-coded). Significant inter-RSN connectivity of the cortical causal network, plotted in a force-directed layout. A clear hub-periphery structure emerged. The dorsal attention network formed a causal pathway from early visual RSNs to multimodal association RSNs. The frontoparietal network was situated in the center of the graph and overall, the most interconnected RSN, receiving directed connections from dorsal attention network, sending directed connections to ventral attention/language, limbic/orbito-affective, and default mode networks, and sharing bidirectional connections with cingulo-opercular network. Percentages on directed connections indicate the proportion of shared connections that were oriented in the statistically preferred direction. B: Cortical surface plot shows the average of nodal in-participation and out-participation coefficient values. Prominent high-participation nodes were apparent in parietal and frontal cortex. C: RSN average participation coefficients. Generally, the posterior multimodal, dorsal attention, frontoparietal, ventral attention, cingulo-opercular, and visual-1 RSNs maintained a high diversity of out-of-RSN connections compared to the rest of the cortex.

The result of this analysis is plotted as a force-directed graph, with connections drawn between RSNs that were found to be significantly connected. Connections are shown as unidirectional if rank-sum testing indicated a preferred direction of connectivity, and connections are shown as bidirectional if rank-sum testing was unable to determine a preferred direction of connections (*Figure 2a*). A clear hub-periphery structure was apparent. We found that visual RSNs 1 and 2 were bidirectionally interconnected, and that visual RSNs projected to the dorsal attention network. The dorsal attention network projected to multimodal association networks (posterior and ventral) and the frontoparietal network. The frontoparietal network was situated in the center of the graph, being the most highly connected RSN and sending information to the ventral attention/language, limbic/orbito-affective, and default mode RSNs, while bidirectionally sharing connections with the cingulo-opercular network. This network connectivity diagram also demonstrated significant bidirectional connectivity between auditory network and the ventral attention/language network, as well as showing strong connectivity between somatomotor and auditory network, which is to be expected since these networks are spatially adjacent and are strongly interconnected (so strongly that they are frequently merged into the same network in the literature; Ji et al., 2019).

We supplemented this analysis by an examination of the average participation coefficient within each RSN. Since we used causal connectivity graphs, we calculated participation coefficient separately using outgoing and incoming connections, resulting in two summary measures per RSN, per subject (hereafter out-part and in-part, respectively). We found that across subjects, the twelve RSNs differed statistically in both out-part and in-part (Friedman tests; both *p* < .001). Post hoc comparisons (Nemenyi test) demonstrated similar patterns of RSN differences for out-part and in-part. The posterior multimodal and dorsal attention RSNs demonstrated the highest participation coefficient values, followed by the frontoparietal, ventral attention, cingulo-opercular, and visual-1 RSNs (all *p <* .001). This analysis of participation coefficients demonstrated that the dorsal attention, posterior multimodal association, frontoparietal, ventral attention, cingulo-opercular, and visual-1 RSNs maintain the greatest diversity of inter-RSN connections in the causal human connectome (*Figure 2b,c*).

### 3.4 The Causal Connectome Has a Heavy-Tailed Centrality Structure

Thus far we have clarified patterns of inter-RSN connectivity in the causal connectome. From here, we examined the most important hubs of the cortical causal connectome using two common centrality metrics: degree centrality, which characterizes highly connected nodes, and betweenness centrality, which characterizes nodes that lie on many shortest paths between other nodes and thus facilitate efficient network communication (Rubinov and Sporns, 2010). If nodes are to be considered hubs based on centrality metrics, the distribution of those metrics should be heavy tailed, with a minority of highly central nodes. We found that the centrality distributions of the obtained causal graphs were indeed heavy-tailed, with most nodes having very low centrality and relatively few nodes having very high causal centrality (*Figure 3a,b,c,d*). Centrality values of the causal connectivity graphs were significantly more heavy-tailed than equally connected random comparison graphs, as confirmed by comparing the skewness of the indegree, outdegree, total degree, and betweenness distributions of these two sets of graphs (rank-sum tests, all *p* < .001). We additionally leveraged this analysis to produce categorical labels of individual brain regions as hubs if their centrality exceeded the 95^th^ percentile of the surrogate distribution. These binary labels almost exclusively designated regions of the lateral and superior parietal cortex as hubs, along with some frontal nodes (*Supplement*).

**Figure 3.**
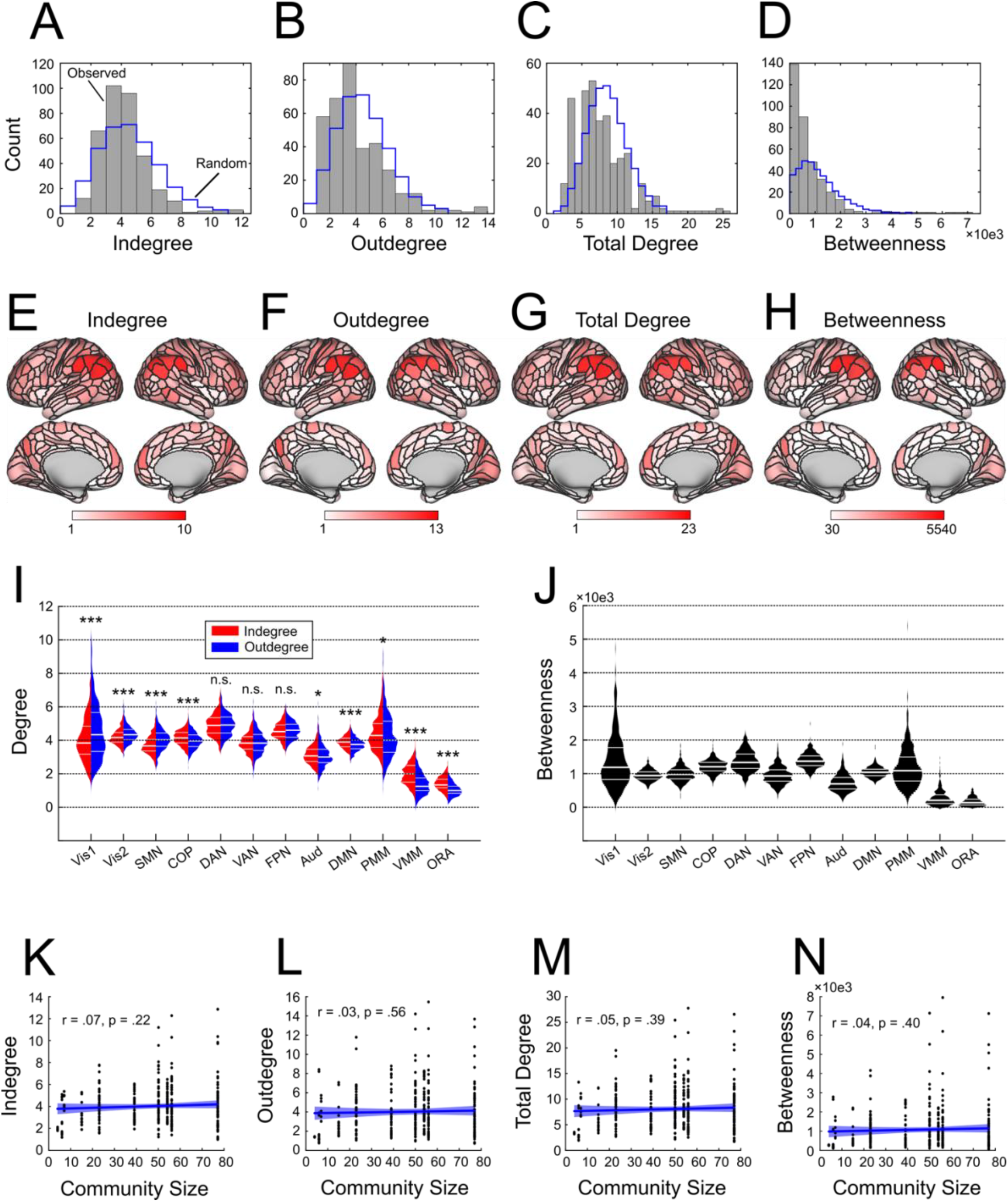
The Most Central Hubs of the Causal Connectome Cluster in Executive and Attentional Networks. **A**,**B**,**C**,**D**: Median nodal centrality across n = 442 subjects, for each of 360 cortical nodes. Blue histogram indicates the distribution of median centralities for 1000 equally connected random graphs. **E**,**F**,**G**,**H**: Centrality values plotted on inflated cortical surfaces to visualize the anatomical locations of highly connected hubs. **I:** Average indegree and outdegree across the 12 RSNs. To determine whether networks primarily send or receive information, for each network we tested for differences between indegree and outdegree. Visual (Vis1 and Vis2) and somatomotor (SMN) networks could be categorized as primarily sending information; cingulo-opercular (COP), auditory (Aud), default mode (DMN), posterior and ventral multimodal (PMM and VMM), and orbito-affective/limbic (ORA) networks could be classified as primarily receiving information, and dorsal attention (DAN), ventral attention/language (VAN), and frontoparietal (FPN) networks equally sent and received information. Overall, the most connected RSNs were the dorsal attention and frontoparietal networks. Asterisks above violin plots indicate significance of the difference between indegree and outdegree for each network (* p < .05, ** p < .01, *** p < .001, n.s. = not significant). **J:** Average betweenness across the 12 RSNs. Overall, frontoparietal network participated in the highest number of short paths, followed by dorsal attention and cingulo-opercular networks. **K**,**L**,**M**,**N:** Control analyses ruled out the possibility that cortical hubs could be explained by the number of parcels in the RSN that each node belongs to. Blue line indicates a least-squares regression fit, and blue shading indicates the 95% confidence interval of the regression.

### 3.5 The Most Central Hubs of the Causal Connectome Cluster in Executive and Attentional Networks

The 12 RSNs differed in average indegree, outdegree, and total degree (Friedman tests, all *p* < .001). Most relevant for the current work, post hoc testing for RSN degree differences (Nemenyi tests) demonstrated that the dorsal attention and frontoparietal networks had significantly higher indegrees, outdegrees, and total degrees than the other 10 RSNs (all *p* < .001). Similarly, comparison of average betweenness centrality across the 12 RSNs (*Figure 3j*) established that frontoparietal nodes participated in the greatest number of efficient paths, followed by dorsal attention nodes; these RSNs had higher betweenness centrality averages than the other 10 RSNs, and frontoparietal had significantly higher betweenness than dorsal attention (all *p* < .001). The cingulo-opercular network had the third highest average betweenness centrality scores, despite having only modest degree centrality, thus suggesting that while cingulo-opercular regions might not be the most highly connected regions in cortex, these regions are nevertheless particularly important for cortical communication. Note that while Power and colleagues (2013) showed that degree-based hubs in Pearson correlation networks are confounded by the size of the functional communities the nodes belong to (i.e, the number of nodes in each RSN), in a critical control analysis we did not find significant correlations between community size and degree or betweenness centrality (*Figure 3k,l,m,n)*, suggesting that our measures of causal centrality cannot be ascribed to the size of the RSNs in our analysis.

### 3.6 Executive and Attentional Networks Equally Send and Receive Connections

Since causal graphs separate incoming and outgoing causal connections, we were additionally able to assess whether each RSN primarily sent or received information. Most RSNs could be characterized as either primarily “senders” (visual, somatomotor), or as primarily “receivers” (cingulo-opercular, auditory, default mode, posterior/ventral multimodal, and limbic/orbito-affective). However, a small number of RSNs were found to send and receive equal numbers of connections (frontoparietal, dorsal attention, ventral attention/language; *Figure 3i*).

### 3.7 Executive and Attentional Hubs are Points of Causal Connectome Vulnerability

Our analyses up to this point have demonstrated that the frontoparietal, dorsal attention, and cingulo-opercular RSNs are highly central hubs in the causal connectome. Based on previous reports, it is likely that the identified central hubs of the causal human connectome are also points of system-level vulnerability to insult. To test this hypothesis, we conducted a series of simulated attacks on the causal connectomes presented in this study. For each RSN, we sequentially deleted nodes in that RSN from each subject’s cortical graph, and measured loss-of-function via percent change in global efficiency (Latora and Marchiori, 2001; Rubinov and Sporns, 2010). *Figure 4a* shows the average of these network-level loss-of-function curves for all 442 subjects. As a summary measure of the impact of nodal targeted attacks on each RSN, we took the average pointwise derivative of the global efficiency loss curve for each subject and RSN (plus random deletion, as a control analysis; *Figure 4b*). Results indicated that the RSN loss functions differed significantly in average pointwise slope (Friedman test, *p* < .001). Post-hoc multiple testing (Nemenyi test) indicated that the frontoparietal network had the steepest loss-of-efficiency function, followed by the dorsal attention network. These RSNs had steeper loss functions than the other 10 RSNs (all *p* < .001). Visual-1, cingulo-opercular, and posterior multimodal network also showed strong efficiency loss effects when lesioned.

**Figure 4.**
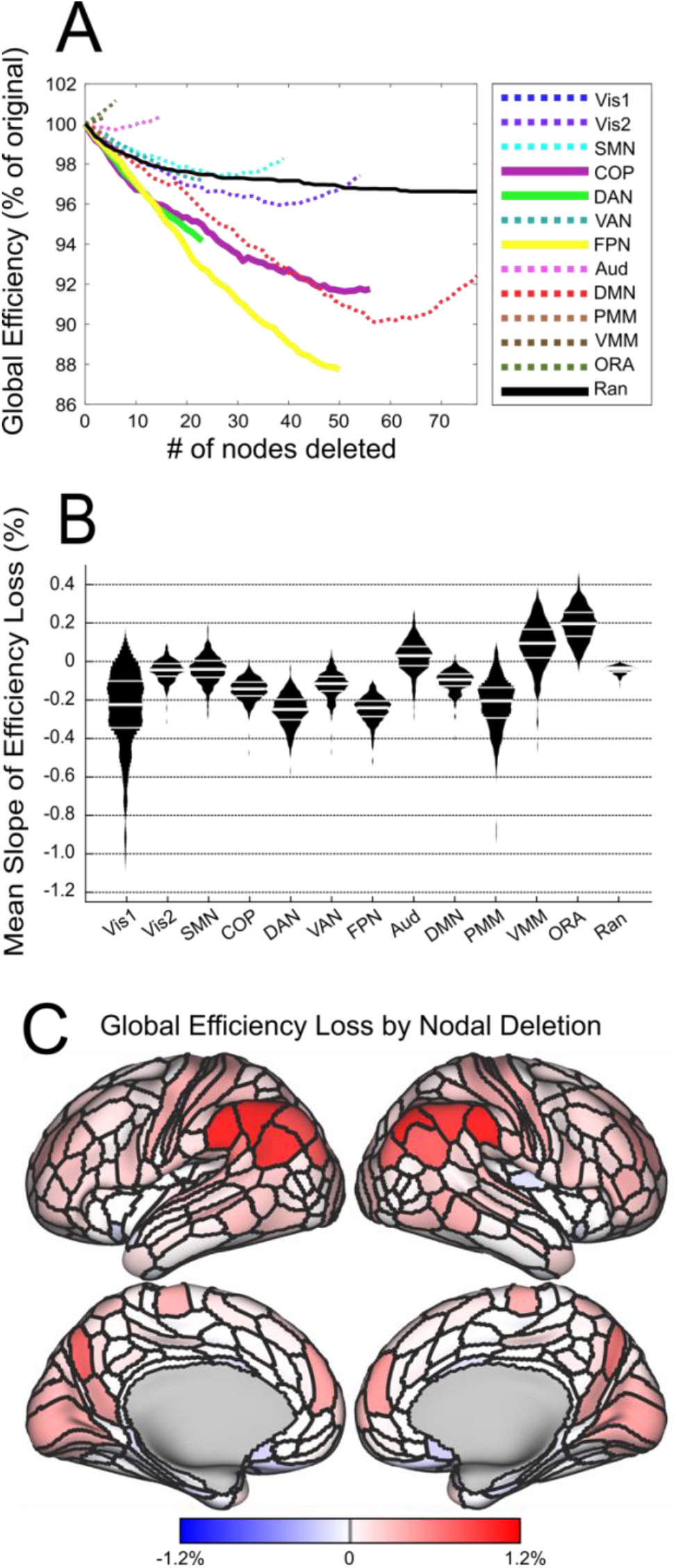
Executive and Attentional Hubs are Points of Causal Network Vulnerability. **A:** Loss of network efficiency following node deletion as a percentage of network global efficiency. The resulting efficiency loss curves are color-coded by RSN. As a visual aid, we plotted RSNs with the most central (i.e. hub-like) connectivity profiles from previous analyses (cingulo-opercular, frontoparietal, dorsal attention) with solid lines, and the remaining (i.e. less central or less hub-like) RSNs are plotted with dotted lines. Additionally, a random attack was carried out (black line) by deleting nodes chosen at random (rather than from a specific RSN). **B:** We calculated the average pointwise slope of the loss curve for each RSN, and compared the average pointwise slopes, thus quantifying how quickly the cortical network loses efficiency when nodes from each RSN are deleted. Violin plots indicate the average slope for the efficiency loss function for each RSN (per subject). **C:** The cortical surface contains nodal values for change in network global efficiency following deletion of individual cortical nodes (that is, the color of each node represents the global efficiency loss when that node is deleted; cold colors indicate gains to global efficiency when the node is deleted).

Counterintuitively, the virtual lesion analysis indicated that for the auditory, ventral multimodal and orbito-affective networks the average global efficiency increased as nodes were deleted. This is likely a result of the very low connectivity of these networks, which have the three lowest degree and betweenness of the RSNs and are among the lowest participation in the connectome as well. This low global connectedness likely means that as nodes are removed the network generally becomes more efficient. In summary, the targeted attack analysis demonstrated that the hub RSNs we previously identified (frontoparietal, dorsal attention, cingulo-opercular) are critical points of vulnerability in cortical efficiency, and that loss-of-function (virtual lesions) in these RSNs impairs global cortical communication efficiency to a greater degree than other RSNs.

### 3.8 Comparison of GANGO Causal Connectivity Graphs with Pearson Correlation Graphs

To examine how the cortical causal human connectome compares to more typical connectivity analyses (Pearson correlation graphs), we ran the presented analyses using two sets of binarized correlation graphs. The first set was proportionally thresholded at a 15% cost (that is, each graph retained the 15% largest positive values). Proportional thresholding was chosen to improve stability of measures over absolute thresholds (Garrison et al., 2015), and the chosen 15% cost is in the middle of an ideal cost range for producing small-world graphs in Pearson correlation networks (Achard and Bullmore, 2007; Bullmore and Bassett, 2011) and as such is among the most typical thresholding procedures in the literature. The second set was thresholded to retain the same number of connections as the subject-specific causal graph, thus matching the density of the causal connectomes exactly. Detailed results of these comparison analyses are presented in the Supplement.

Overall, this comparison suggests that Pearson correlation graphs emphasize the importance of sensory regions and motor cortex as cortical hubs, while causal connectivity graphs instead emphasize higher cognitive regions, particularly frontoparietal and cingulo-opercular networks (which did not exhibit hub-like connectivity in any analysis for Pearson correlation graphs). We also found that the sparser set of correlation graphs were very poorly connected – on average, over 40% of nodes were completely disconnected at this threshold. At this 2.25% (average) density, we also found that graphs were significantly *less* hub-like than surrogate random graphs, suggesting that centrality-based hubs break down at this level of sparsity in Pearson correlation graphs.

Furthermore, in 15% density Pearson correlation graphs we found that both degree and betweenness centrality were highly confounded by the size of the RSN nodes belonged to, unlike in the causal graphs. This was not the case for the sparser correlation graphs. These differences between Pearson correlation and causal connectomes were somewhat attenuated when using participation coefficient as a measure. However, in the sparse 2.25% density correlation graphs, we were unable to estimate participation coefficient for the two least-connected networks (ventral multimodal, limbic/orbito-affective) due to extremely low levels of connectivity (zero in nearly all subjects). We also observed that applying the virtual lesion analysis to the 15% density Pearson correlation graph resulted in many RSNs increasing global efficiency when deleted. This effect was magnified for the sparser 2.25% density correlation graphs. This is likely due to the much greater incidence of completely unconnected nodes in thresholded Pearson correlation graphs, as opposed to the well-connected causal graphs we generated using the GANGO method. Note that unconnected nodes have this effect because the global efficiency of a node is calculated as the inverse shortest path (number of edges) from that node to each other node in the network (Latora and Marchiori, 2001). The global efficiency of the network is then the average of all nodal global efficiencies. An unconnected node is defined to have an infinite path length and a global efficiency of zero (Rubinov and Sporns, 2010) – thus, these unconnected nodes reduce the average global efficiency (connected nodes will always have global efficiency >0). It follows then, that the deletion of an unconnected node removes a zero from being averaged into the global efficiency score, improving the average network efficiency.

## 4 Discussion

Functional connectivity analyses of the human cortex typically use undirected connectivity estimates, derived by computing Pearson correlation coefficients between the time series of the hemodynamic signals of individual brain regions (nodes). The need to extend functional connectivity analyses to causal connectivity is recognized (Reid et al., 2019; Smith, 2012), but data-driven methods for calculating high-dimensional causal graphs within single subjects have not been thoroughly tested yet. Here, we present an examination of the causal connectivity patterns of the human cortex using a two-stage causal discovery machine learning approach. Two-stage causal discovery methods break the graph creation process into separate adjacency search and orientation phases. Ramsey et al. (2014) and Sanchez-Romero et al. (2019) both used the PC algorithm (Spirtes et al., 2001) for the adjacency search, and demonstrated that non-Gaussian pairwise likelihood measures (particularly skew-based measures) could accurately identify correct edge orientations. We demonstrate the utility of a scalable version of these two-step causal discovery algorithms, which we call GANGO for convenience. Scalability is achieved by substituting the PC adjacency search with FGES (Fast Greedy Equivalence Search; Ramsey et al., 2017), itself a parallelized version of Greedy Equivalence Search (Chickering, 2002), which was shown to produce high precision for adjacency search in Smith et al. (2011).

The causal connectomes produced by the GANGO approach were quite different from those produced by the more typical Pearson correlations - despite very low density, these graphs were fully connected in nearly all cases. For this initial justification of the GANGO approach, we used the standard BIC score (penalty discount = 1) to penalize the connectivity density of the produced graphs; notably, the FGES method can be parameterized to produce sparser graphs with penalty discounts >1, and to produce denser graphs with penalty discounts <1. As such, future applications of the GANGO framework might capitalize on this flexibility to produce graphs with the desired density for the research question under investigation. Furthermore, GANGO networks did not exhibit any relationship between RSN size (number of nodes) and degree centrality, unlike standard Pearson correlation-based graphs. This dependency arises in correlation-based graphs whenever the graph exhibits a modular community structure, as explained in Power et al. (2013). Specifically, in cases where graphs exhibit community structure, a node in a larger community would have a higher chance to form more connections simply because connections are more common within communities than between them. However, many of these connections will be indirect, and nodal correlations within a community will be high due to indirect connections. Bayes net methods, on the other hand, enforce sparsity wherever possible via a Markovian screening-off property and retain only direct connections while eliminating indirect connections (Spirtes et al., 2001). Thus, our results provide support for the viability of the GANGO approach for providing unbiased centrality measures from resting-state fMRI data.

Prominent hubs of the causal connectome overlap many regions previously identified by resting-state fMRI (Achard et al., 2006; Buckner et al., 2009; Tomasi and Volkow, 2011; van den Heuvel and Sporns, 2013; Zuo et al., 2012), with the GANGO method reliably recovering these network properties when applied on a single subject level. Importantly, control analyses indicated that nodal hub metrics (degree, betweenness centrality) were unconfounded by the size of the RSN that nodes belonged to. In contrast, degree and betweenness centrality were strongly confounded by RSN size for thresholded correlation graphs (Supplement). Overall, we found prominent causal connectivity hubs and points of vulnerability of the causal connectome in dorsal attention network (DAN), frontoparietal network (FPN) and cingulo-opercular network (COP), with each of these hub networks showing distinctly different connectivity profiles.

The dorsal attention network (DAN) exhibited theoretically interesting properties that contributed to its high level of connectivity. In our analysis, DAN had among the greatest diversity of connections with other RSNs (measured using participation coefficient), as well as having overall high connectivity (measured using degree centrality) and participating in many efficient paths (measured using betweenness centrality). Our analysis of the inter-RSN connectivity structure of the cortical causal connectome revealed that DAN owed its high centrality to its role in receiving information from visual networks, processing that information, and then transmitting information to multimodal association networks (posterior/ventral multimodal association networks, FPN). This is in line with long-standing evidence that DAN plays an important role in top-down visual selective attention (Corbetta and Shulman, 2002; Vossel et al., 2014). While we found that causal connections usually progressed from visual to dorsal attention networks, a large proportion of these connections still progressed in the opposite direction as well. Thus, our results support a role of DAN in top-down control over visual systems as well, providing further evidence that the dorsal attention network supports both bottom-up sensory integration and top-down attentional control (Long and Kuhl, 2018).

The cingulo-opercular network (COP) was found to mediate many efficient paths in the cortex (betweenness) and shared a large diversity of inter-RSN connections (participation coefficient) but did not have particularly high connectivity (degree). COP has a role in maintaining task sets, initiating goal-directed behaviors, and consolidating motor programs (Dosenbach et al., 2008, 2006; Fair et al., 2007; Newbold et al., 2021). In our consensus network structure, we found that COP received significant connections from the somatomotor network, sent significant connections to the orbito-affective (reward) network, and was bidirectionally connected to FPN. The uncovered functional connectivity between the reward networks and COP is in line with evidence that COP has a role in coordinating the response of brain reward-related regions (Huckins et al., 2019), and the connectivity between COP and somatomotor networks corroborates evidence that COP plays a role in consolidation, planning, and plasticity of motor regions (Newbold et al., 2021). Connectivity between COP and somatomotor networks also increases through development and is linked to the development of improved cognitive control (Marek et al., 2015). Finally, COP was found to be tightly bidirectionally connected with frontoparietal network, echoing evidence that these RSNs work together as dual cognitive control networks (Dosenbach et al., 2008; Fair et al., 2007; Gratton et al., 2018).

Across all analyses, we found a critical role of frontoparietal executive network (FPN) connectivity. This represents an important point of agreement between our causal network results and recent advances in understanding the role of the FPN in overall function, as well as an important result that we did not find with traditional Pearson correlation graphs (Supplement). Due to its central position, FPN shared the greatest diversity of inter-RSN connectivity in the consensus graph, including significant received connections from DAN, significant sent connections to the ventral attention, orbito-affective, and default mode networks, and bidirectional connections with COP. Previous studies demonstrate that FPN flexibly shifts its connectivity patterns to fulfill task demands, while still retaining high correlations with its resting-state connectivity (Cocuzza et al., 2020; Cole et al., 2013; Crittenden et al., 2016), and another recent study demonstrated that resting-state network connectivity in FPN predicts transfer of task-relevant information through distributed brain circuitry (Ito et al., 2017). Overall, our finding that FPN nodes are, on average, the most highly central nodes in the causal human connectome is consistent with a theoretical role of FPN as a flexible executive coordinator of overall brain function (Assem et al., 2020; Dosenbach et al., 2006; Duncan, 2010; Fair et al., 2007; Marek and Dosenbach, 2018).

This is also consistent with recent control systems perspectives on brain connectivity, which have suggested that FPN has a particular role in shifting brain network configuration into difficult-to-reach cognitive states (Gu et al., 2015). The causal connectivity from FPN to default mode network (DMN) deserves particular attention. These RSNs are typically anticorrelated during cognition (FPN activity increases and DMN activity decreases), a finding that replicates in experimental paradigms including attention (Hellyer et al., 2014), working memory (Kelly et al., 2008; Murphy et al., 2020), and cognitive reasoning (Hearne et al., 2015). We found that the connectivity between FPN and DMN preferentially flows from FPN to default mode network, suggesting that FPN might have an executive role in “turning off” or inhibiting the DMN in response to the need for control. The revealed causal hub role of the FPN is among the most important contributions of this study, as typically thresholded Pearson correlation graphs do not show hub-like connectivity in FPN (Supplement), despite ample theoretical and experimental evidence that the frontoparietal network is critical for overall organization and control of the connectome.

Our virtual lesion analysis of the causal human connectome suggests a potential application of the GANGO method to understand brain impairments in psychiatric disorders. A previous analysis (Crossley et al., 2014) demonstrated that virtual lesions to the hubs of the human connectome impair network global efficiency more than virtual lesions of non-hub brain nodes. The authors then followed this with a meta-analysis demonstrating that the hubs of the human connectome were more likely to contain grey matter lesions than non-hub regions across nine different disorders, including schizophrenia and Alzheimer’s disease. Many psychiatric and neurological disorders are associated with reduced brain global efficiency, including prenatal alcohol exposure (Wozniak et al., 2013) and fetal alcohol syndrome (Rodriguez et al., 2021), schizophrenia (Hummer et al., 2020), ADHD (Wang et al., 2020; Wang et al., 2021), generalized anxiety disorder (Guo et al., 2021), heavy smoking (Lin et al., 2015), and major depressive disorder (Meng et al., 2013; Wang et al., 2017; Zhi et al., 2018), among others. Additionally, prefrontal tDCS for alcohol use disorder increased brain network global efficiency (Holla et al., 2020), suggesting that normalizing network global efficiency might contribute to improved treatment outcomes from neuromodulation therapy. Furthermore, mindfulness-based cognitive therapy in mood-dysregulated adolescents resulted in an increase in brain global efficiency, especially within frontoparietal and cingulo-opercular networks (Qin et al., 2021). GANGO causal connectomes, but not standard Pearson correlation connectomes, emphasize hub connectivity in frontoparietal and cingulo-opercular networks, as well as suggesting that these networks are points of vulnerability with regards to their impact on global network integration when lesioned. Thus, the causal human connectome might explain the high incidence of frontoparietal dysfunction and global efficiency reduction in patient groups, as well as predicting the therapeutic effects of frontoparietal stimulation in various psychiatric dysfunctions. As such, the GANGO framework might provide a powerful new tool to understand, predict, and ultimately treat brain network dysfunction in psychiatry.

The current investigation delivers a powerful new framework for quickly computing (∼30 s per connectome on a personal laptop with six cores and 32 Gb Ram) high-dimensional causal connectivity graphs from observed brain data as well as providing important insight into the hub structure of the causal human connectome, but it is not without limitations. One potential limitation lies in the use of a relatively coarse (n = 12) RSN partition for summarizing cortical hubs (Ji et al., 2019). However, the use of a published RSN partition facilitates interpretation of results, as the results of higher-dimensional (e.g., ICA-based) RSN partitions are often difficult to interpret and require abstraction via multidimensional statistics to summarize. In the Supplement we report the consensus network structure of these same data using a larger number of RSNs [the 22 neuroanatomical regions reported in the supplement of (Glasser et al., 2016a)]. The consensus structure of connectivity between these 22 regions also shows an orderly progression of information from visual sensory regions to the dorsal and ventral visual streams, through the parietal association cortex, and into the motor and prefrontal cortex. The organization of this more granular causal network also aligns well with recent perspectives on the hierarchical organization of the prefrontal cortex (Badre and Nee, 2018). An additional limitation of these results is that our method for calculating causal connectivity is unable to discover two-cycles (direct feedback cycles, where A->B and B->A). While two methods have been proposed for fMRI connectivity that are theoretically capable of recovering two-cycles (Sanchez-Romero et al., 2019), these methods have never been used in an applied research context, and often perform worse than the skew-based orientation method we use (Sanchez-Romero et al., 2019). Notably, the RSkew orientation method we adopt for the GANGO framework can discover 3-cycle or greater feedback loops, so only direct feedback loops remain unmeasured. Nevertheless, as methods for more accurately assessing feedback cycles from fMRI are developed, the framework we implement in the current investigation could be expanded to include such methods. Finally, we note that, while every single subject in this study had significantly skewed BOLD distributions (with reference to random Gaussian data), this is not guaranteed to be the case in all datasets. As some common preprocessing steps (aggressive temporal filtering) can introduce Gaussian trends into BOLD data, we recommend that application of the GANGO framework should only follow careful examination of BOLD skewness, to ensure that the assumptions of the methods are adequately met.

## 5 Conclusion

Using a causal discovery machine learning framework, we demonstrate that the most centrally connected hubs of the cortical connectome are situated in the frontoparietal, dorsal attention and cingulo-opercular networks. In particular, the causal human connectome highlights high connectivity of the frontoparietal network with all other higher cognitive RSNs. The discovered hub role of the frontoparietal network in the causal human connectome is especially attractive, as brain-based therapies for psychiatric conditions typically impact or directly stimulate nodes in the frontoparietal network (Belsher et al., 2021; Ferrarelli and Phillips, 2021; Fitzgerald, 2021; Song et al., 2019; Voigt et al., 2021; Zhang et al., 2021; Zilverstand et al., 2017, 2016). Previously, we even demonstrated that connectivity in the frontoparietal network has downstream causal effects on the severity of alcohol use disorder (Rawls et al., 2021). As it is applied on a single-subject basis, the GANGO method could potentially even enable individualized causal connectivity-based neuromodulation targeting. Thus, the current study sets the stage for future applications of data-driven causal connectivity applications in psychiatry.

## Acknowledgements

Data were provided by the Human Connectome Project, WU-Minn Consortium (Principal Investigators: David Van Essen and Kamil Ugurbil; 1U54MH091657) funded by the 16 NIH Institutes and Centers that support the NIH Blueprint for Neuroscience Research; and by the McDonnell Center for Systems Neuroscience at Washington University. ER is supported by a postdoctoral training grant from the National Institutes of Mental Health (T32-MH115866). The content is solely the responsibility of the authors and does not necessarily represent the official views of the National Institutes of Mental Health.

## Author Contributions

**ER:** Conceptualization, Methodology, Software, Validation, Formal Analysis, Writing – Original Draft, Writing – Review and Editing, Visualization, Funding Acquisition. **EK:** Conceptualization, Methodology, Validation, Writing – Review and Editing, Supervision. **BAM:** Validation, Writing – Review and Editing. **SM:** Validation, Writing – Review and Editing. **AZ:** Conceptualization, Methodology, Validation, Resources, Writing – Review and Editing, Supervision, Project Administration, Funding Acquisition.

## Data and Code Availability

Data were provided by the Human Connectome Project, WU-Minn Consortium, and are publicly available at https://db.humanconnectome.org/data/projects/HCP_1200. Fast greedy equivalence search (FGES) was conducted using causal-cmd software, available at https://bd2kccd.github.io/docs/causal-cmd/. Pairwise likelihood ratios for edge orientations were computed using MATLAB code provided by A. Hyvärinen and S. M. Smith, available from the authors at https://www.cs.helsinki.fi/u/ahyvarin/code/pwcausal/. All graph theory metrics were computed using the Brain Connectivity Toolbox (BCT), available at https://sites.google.com/site/bctnet/.

