## Supplement for "The Resting-State Causal Human Connectome is Characterized by Hub Connectivity of Executive and Attentional Networks"

### Supplemental Results 1.1: Illustration of Significant Nodal Hubs of the Causal Human Connectome

As described in Sections 2.5 (Methods) and 3.4 (Results) of the accompanying manuscript, we leveraged our permutation analysis of causal connectome centrality distributions to find significant nodal hubs of the cortex (*Figure S1*). A nodal hub was defined as significant if that node's centrality passed the 95th percentile of the random surrogate distribution. For indegree, outdegree, and betweenness, this analysis identified only hubs in superior and lateral parietal cortex, primarily assigned to dorsal attention and frontoparietal networks. For total degree, this analysis also identified a hub in lateral prefrontal cortex (part of FPN) and a hub in medial frontal cortex (part of default mode network).

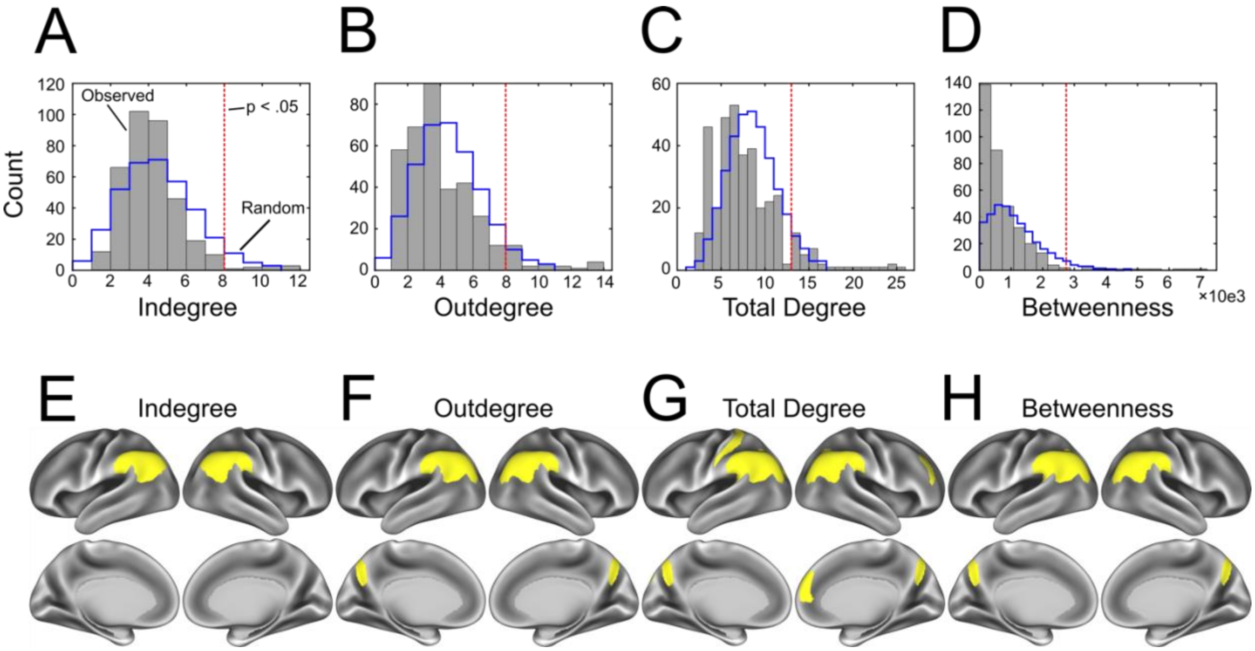


*Figure S1. We applied our permutation-based analysis of centrality distributions to identify significant hubs in the causal human connectome.*
*A, B, C, D: Same as Figure 2a,b,c,d in the accompanying manuscript, but with the addition of empirical cutoffs for defining regions as hubs or non-hubs, for each main centrality metric we consider.*
*E,F,G,H: We plotted regions that were found to have significantly hub-like connectivity on an inflated cortical surface, to enable inspection of nodal hub assignments in the causal connectome.*

**Supplemental Results 1.2: Comparison of Effective Cortical Network Results to Correlation Graphs**

To examine how the cortical causal connectome compares to a more typical connectome derived from Pearson correlations, we ran all the presented analyses also using binarized correlation graphs, thresholded using two different cost levels. For the first of these comparisons, we proportionally thresholded the Pearson correlation graphs at 15% cost (i.e., each graph retained the 15% largest positive values) based on the most typical thresholding procedures from the literature (Achard & Bullmore, 2007; Bullmore & Bassett, 2011). For the second set of comparison graphs, we retained the same number of edges as each subject's causal connectome (mean density = 2.25%, varied per person). While this second threshold is unusually sparse for correlation graphs and is outside of the range of values that produce small-world attributes in correlation networks (Achard & Bullmore, 2007), we considered this threshold necessary for comparison to the equally dense causal graphs.

Our analysis of the inter-network connectivity structure of the correlation graphs (15%: *Figure S2a*, 2.25%: *Figure S3a*) showed some of the same in-network connectivity patterns as the effective network analysis, but with a less pronounced hub structure and without directionality. For example, both levels of correlation networks showed significant connectivity between visual1 (Vis1) and visual2 (Vis2) networks, and that visual networks and cortical association networks were bridged via connections passing through the dorsal attention network (DAN). In the 15% correlation graphs, connectivity between higher cognitive networks was less interpretable than that in the causal connectome; for example, this analysis did not reveal known strong links between cingulo-opercular (COP) and frontoparietal (FPN) networks (Fair et al., 2007; Gratton et al., 2018), and did not suggest any important role of higher cognitive networks as cortical hubs. The sparse (2.25%) correlation graphs unexpectedly produced maps of inter-network connectivity that were more like the causal connectome – at this sparsity, we recovered a central role of FPN in the force-directed network graph, as well as uncovering known connectivity between COP and FPN. However, likely due to high levels of disconnection in these sparse graphs, we found that ventral multimodal (VMM) and limbic/orbito-affective were not significantly connected to any other RSNs (in the 15% connectomes, ORA was similarly disconnected).

In line with prior reports that participation coefficient might be a more suitable metric to define hubs in correlation graphs as compared to other measures such as degree, the average network participation coefficients derived from the Pearson correlation graphs were relatively similar to those presented for the effective connectivity graphs in the main manuscript (15%: *Figure S2b,c*, 2.25%: *Figure S3b,c*). Specifically, the 12 networks differed significantly in participation coefficient (Friedman $p < .001$) at both threshold levels. For 15% density graphs, post-hoc testing (Nemenyi test) showed that posterior multimodal network had higher participation coefficient than the other 11 networks ($p < .001$), DAN and Vis1 networks had higher participation coefficients than the other 9 networks ($p < .001$) but did not differ from each other, and Vis2, somatomotor (SMN), and ventral attention/language network (VAN) had higher participation coefficients than the other six networks ($p < .001$) but did not differ from each other. For the sparse (~2.25%) density correlation graphs, DAN had higher participation coefficient than the other 11 networks ($p < .001$), PMM and Vis1 networks had higher participation coefficients than the other 9 networks ($p < .001$) but did not differ from each other, and Vis2, SMN, COP, FPN, and VAN had higher participation coefficients than the other five networks ($p < .001$) but did not differ from each other. As such, both thresholding levels generated nearly identical participation coefficient results (at least at the RSN level),

We continued to investigate the centrality distributions of the correlation graphs, to establish whether these graphs are compatible with a hub structure containing a few highly connected nodes. The degree histogram of the 15% density correlation graphs was not heavy tailed, but rather was closer to a uniform distribution with a notable peak at zero (i.e., many nodes were disconnected; *Figure S2d*). The degree histogram of the sparse 2.25% graphs showed an even greater peak at zero (i.e., almost all nodes were disconnected) with an exponential decay (*Figure S3d*). The distribution of betweenness centrality values in both the 2.25% graphs and the 15% cost graphs were like that obtained for the causal connectivity graphs, however, with many low-centrality nodes and a small number of high-centrality nodes (*Figure S2e, Figure S3e*). Importantly, for the 2.25% cost graphs, the tail of the surrogate distribution was significantly *longer* than that of the observed graphs, and thus the sparse correlation graphs are significantly *less* hub-like than random graphs. As such, we do not analyze betweenness centrality further for the 2.25% sparsity graphs.

In the 15% cost graphs, high-degree nodes were widely spread over the visual/occipital cortex, parietal cortex, and some frontal nodes (*Figure S2d*). This distribution was similar for the sparser 2.25% graphs (*Figure S3d*). For the 15% density graphs, the 12 networks differed markedly in average degree (Friedman $p < .001$), with post-hoc testing (Nemenyi test) demonstrating that the highest-degree networks were visual (1 & 2), posterior multimodal (PMM), and DAN, followed by SMN. For the sparser 2.25% (average) density graphs, the 12 networks differed markedly in average degree (Friedman $p < .001$), with post-hoc testing (Nemenyi test) demonstrating that the highest-degree networks were Vis2, DAN, and Vis1, followed by SMN and PMM. The frontoparietal network notably had a low average degree in both the 2.25% and the 15% correlation graphs, at odds with the hub role of the frontoparietal network that we found in effective connectivity graphs (*Figure S2f*).

For the 15% cost correlation graphs, the 12 networks also differed significantly in betweenness centrality (Friedman $p < .001$), and post-hoc testing (Nemenyi test) demonstrated a similar pattern as that found for degree. Vis1 and PMM had the highest betweenness centrality, followed by DAN (*Figure S2g*). Unlike the analysis of effective connectivity, which demonstrated that frontoparietal network had the highest betweenness centrality of the 12 RSNs, in correlation graphs the frontoparietal network had only average centrality (ranked 5th out of 12). We did not run this comparison for the sparser 2.25% correlation graphs, since these graphs did not exhibit any hub-like connectivity using betweenness. Perhaps most importantly, in a control analysis we demonstrated that both centrality measures (degree and betweenness) were heavily confounded by the size of the functional community a node belonged to (i.e., the number of parcels in the RSN; *Figure S2h*). Intriguingly, however, this analysis found significantly negative correlations between centrality and community size, unlike Power et al. (2013), who reported positive correlations between degree and community size. This also was only the case for the 15% density correlation graphs; degree was unconfounded by community size for the 2.25% density graphs (possibly due to the inflated number of unconnected nodes, over 40% of nodes on average). Nevertheless, our results do confirm that in correlation networks, at least at the cost threshold usually applied in the literature, centrality metrics are confounded by the size of the RSN nodes belong to. Finally, the virtual lesion analysis also indicated that the effect of node deletion on overall network efficiency differed between the 12 RSNs for both sets of thresholded Pearson correlation graphs (Friedman $p < .001$). Post-hoc testing (Nemenyi test) demonstrated that for both sets of correlation graphs, the most impactful networks were visual-1, posterior multimodal, dorsal attention, and visual-2 networks (*Figure S2i,j,k*). Results indicated that random deletion impacted overall efficiency more than targeted deletion of the remaining networks. This result again demonstrates that correlation graphs tend to emphasize connectivity of brain visual pathways, while the causal connectomes presented in the main manuscript instead accentuate the importance of higher cognitive networks (the frontoparietal network). Counterintuitively, the virtual lesion analysis indicated that for some networks (most notably default mode network [DMN]), the average network global efficiency increased monotonically for nodal deletion. A likely reason for this can be found in examination of participation coefficient values: DMN nodes can be seen to primarily connect within the DMN, generally having very low participation coefficient values. This low global connectedness likely means that as these nodes are removed, the network generally becomes more efficient, since other networks are more globally connected than the DMN. These effects were magnified for the sparser 2.25% density graphs, likely due to the poor overall connectedness of these graphs.

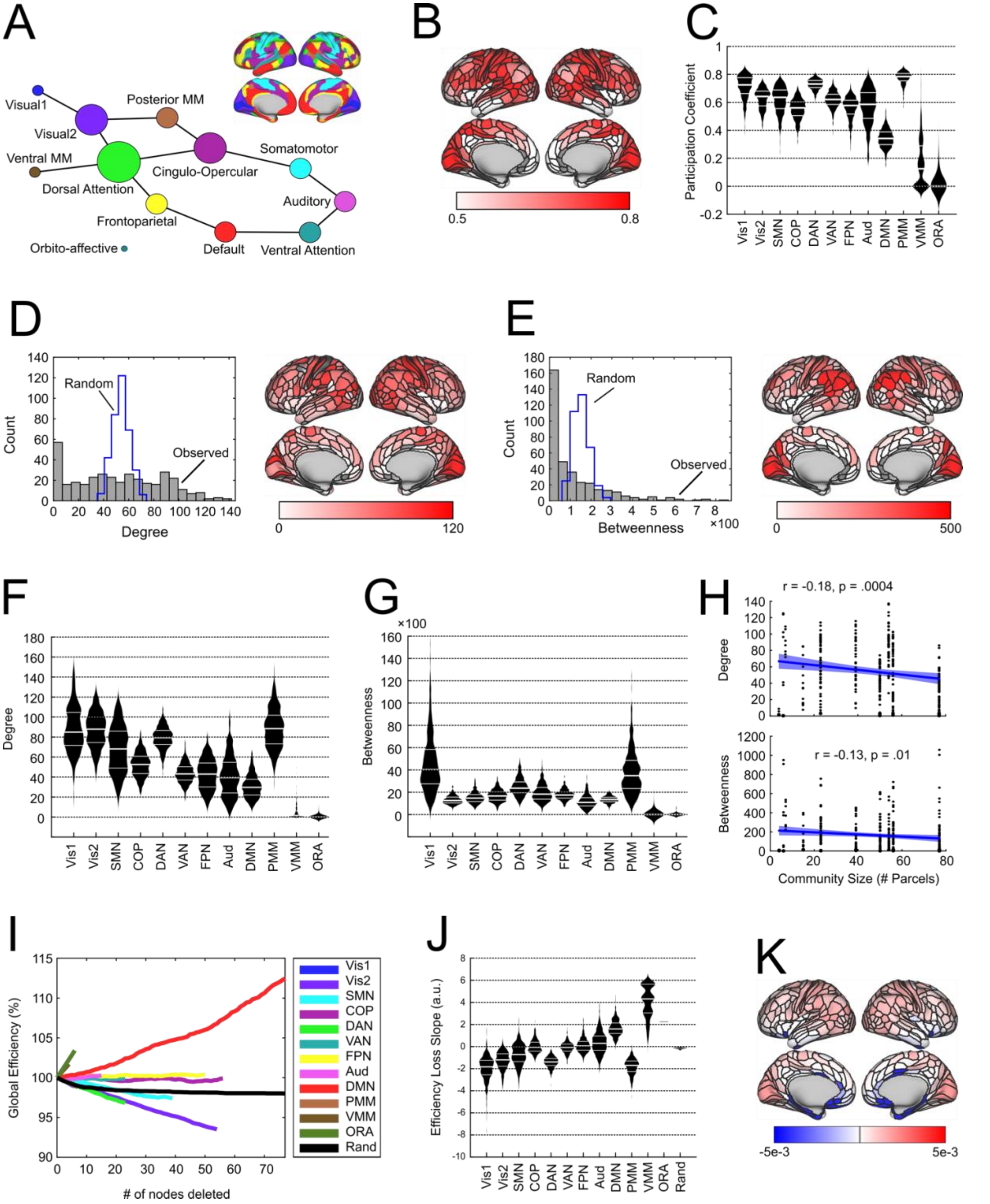


*Figure S2. We ran all of the previously described analyses on correlation graphs, thresholded proportionally at a cost of 15%, to compare the effective connectivity results to the more typical correlation graphs.*

***A**: Significant inter-network connectivity structure of the cortical correlation network. For each of the 12 networks, we established whether pairs of networks shared a significant number of connections compared to expected random connectivity.*

***B.*** *Distribution of participation coefficient values across cortical nodes.*
***C.*** *Network average participation coefficient values.*
***D****: Histogram indicates the median nodal degree across n = 442 subjects, for each of 360 cortical nodes. Blue stairs indicate the median nodal degree across 1000 random graphs. Inflated cortical surface shows the distribution of degree across cortical nodes.*
***E****: Histogram indicates the median betweenness centrality across n = 442 subjects, for each of 360 cortical nodes. Blue stairs indicate the median betweenness centrality across 1000 random graphs. Inflated cortical surface shows the distribution of betweenness centrality across cortical nodes.*
***F,G****: Network average degree (F) and betweenness centrality (G).*
***H:*** *Both degree and betweenness centrality were highly confounded by the size of the RSN (number of parcels comprising the network), although with the opposite sign presented by Power et al. (2013).*
***I:*** *We sequentially deleted each node comprising each of the 12 RSNs we analyzed and recorded the loss of network efficiency following node deletion as a percentage of network global efficiency. The resulting efficiency loss lines are shown, color-coded by RSN. Additionally, a random attack was carried out (black line) by deleting an equivalent number of nodes chosen at random (rather than from a specific RSN).*
***J:*** *We calculated the slope of the loss line (via linear regression) for each network, and compared the slopes, thus quantifying how quickly the cortical network loses efficiency when nodes from each network are deleted.*
***K:*** *Loss-of-efficiency for deletion of each cortical node in the network, plotted on the inflated cortical surface.*

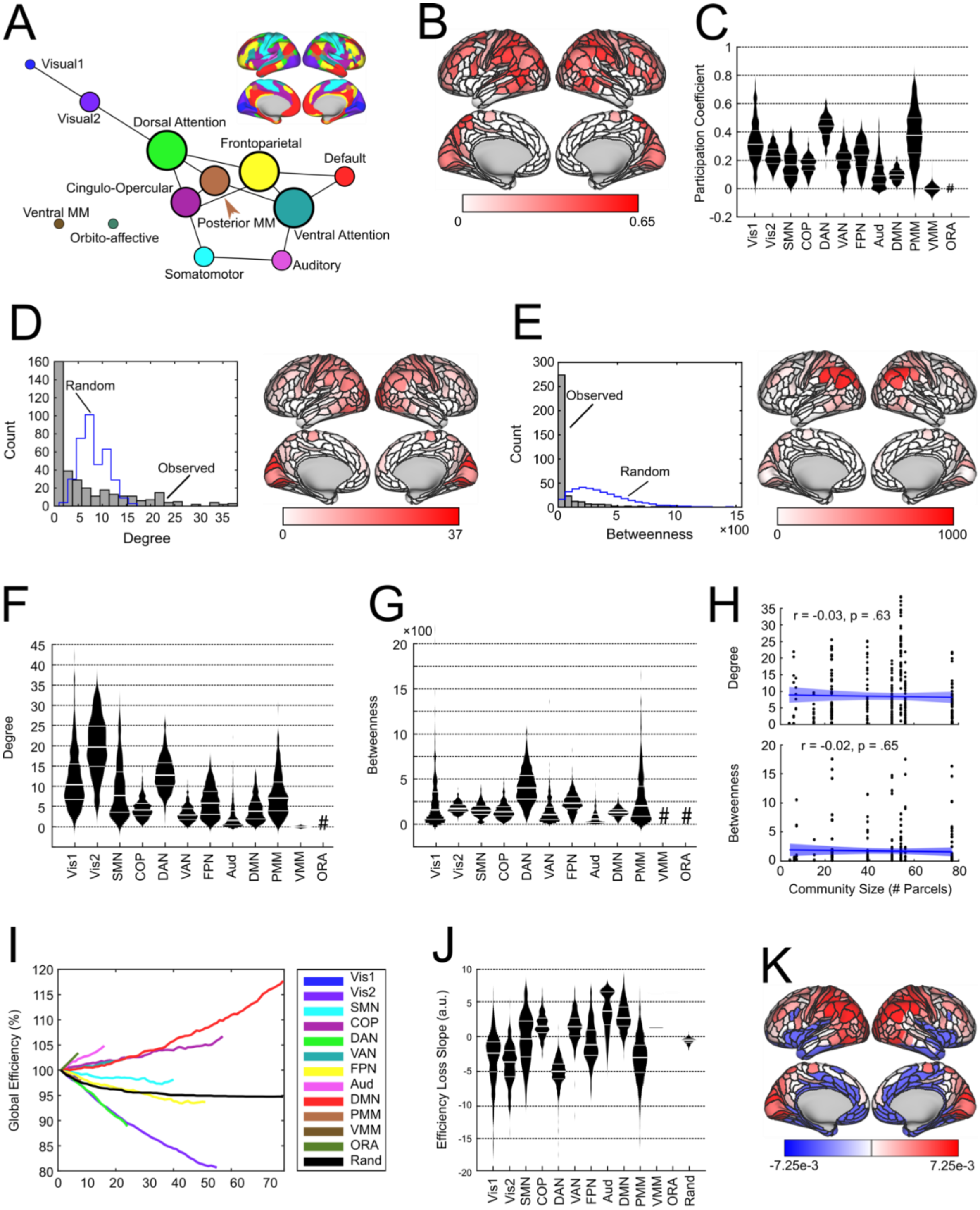


*Figure S3. We ran all of the previously described analyses on correlation graphs, thresholded to have the exact density of the causal connectomes presented in the accompanying main manuscript, to compare the effective connectivity results to equally dense correlation graphs.*

***A**: Significant inter-network connectivity structure of the cortical correlation network. For each of the 12 networks, we established whether pairs of networks shared a significant number of connections compared to expected random connectivity.*

***B.** Distribution of participation coefficient values across cortical nodes.*

***C.*** *Network average participation coefficient values.*
***D****: Histogram indicates the median nodal degree across n = 442 subjects, for each of 360 cortical nodes. Blue stairs indicate the median nodal degree across 1000 random graphs. Inflated cortical surface shows the distribution of degree across cortical nodes.*
***E****: Histogram indicates the median betweenness centrality across n = 442 subjects, for each of 360 cortical nodes. Blue stairs indicate the median betweenness centrality across 1000 random graphs. Inflated cortical surface shows the distribution of betweenness centrality across cortical nodes.*
***F,G****: Network average degree (F) and betweenness centrality (G).*
***H:*** *At this low density, neither degree nor betweenness centrality were confounded by the size of the RSN (number of parcels comprising the network).*
***I:*** *We sequentially deleted each node comprising each of the 12 RSNs we analyzed and recorded the loss of network efficiency following node deletion as a percentage of network global efficiency. The resulting efficiency loss lines are shown, color-coded by RSN. Additionally, a random attack was carried out (black line) by deleting an equivalent number of nodes chosen at random (rather than from a specific RSN).*
***J:*** *We calculated the average slope of the loss line via pointwise derivative for each network, and compared the slopes, thus quantifying how quickly the cortical network loses efficiency when nodes from each network are deleted.*
***K:*** *Loss-of-efficiency for deletion of each cortical node in the network, plotted on the inflated cortical surface.*

**Supplemental Results 1.3: Examination of Causal Connectivity in a More Granular Set of Communities**

Given the results presented in the main manuscript, one might wonder whether the GANGO effective connectivity method is able to recover inter-modular connectivity for a more fine-grained set of networks, compared to the 12 networks we present. Indeed, a potential limitation of the main results lies in the use of a relatively coarse (n = 12) network partition for summarizing cortical hubs (Ji et al., 2019), despite that the use of a published network partition facilitates interpretation of results. We ran the analysis described in Section 4.4 (Network Connectivity Statistics) using the 22 neuroanatomically defined regions reported in the supplement of Glasser et al., (2016) rather than the 12 networks reported in (Ji et al., 2019), to obtain a diagram of the significant inter-network effective connectivity with a greater degree of granularity (*Figure S4*). The consensus structure of connectivity between these 22 regions shows an orderly progression of information from visual sensory regions to the dorsal and ventral visual streams, through parietal association cortex, and into motor and prefrontal cortex. Results of the connectivity structure within frontal cortex also parallel evidence of a processing hierarchy within prefrontal cortex (Badre & Nee, 2018).

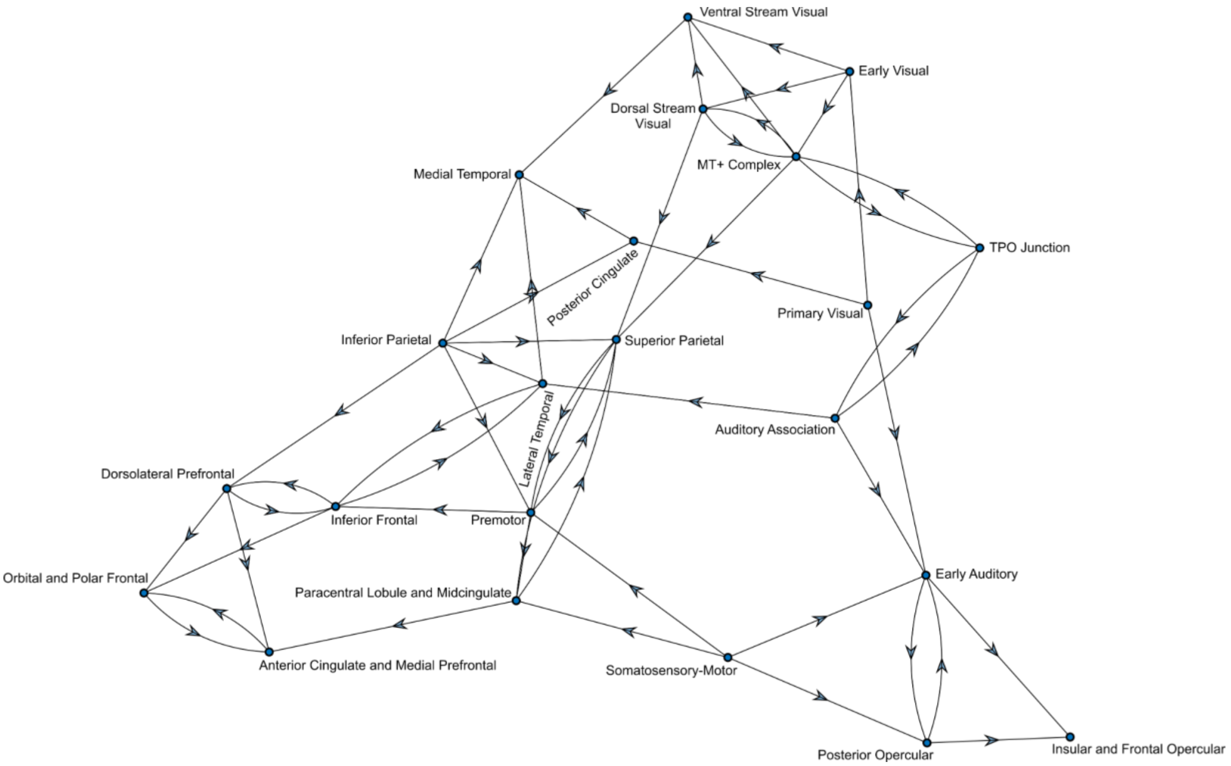


*Figure S4. We applied our inter-network connectivity analysis to a more granular set of 22 neuroanatomical regions reported in the supplement of (Glasser et al., 2016) in place of the 12 networks we used in the main body of the manuscript.*
